# Impact of oxygen fluctuation and consumption on the dynamics of a reduced go-or-grow model for tumor cell invasion

**DOI:** 10.64898/2026.10.02.756316

**Authors:** Gopinath Sadhu, Mohit Kumar Jolly, Philip K Maini

## Abstract

We propose a minimal mathematical framework to describe the go-or-grow dynamics of tumor cells comprising two phenotypically distinct populations. One population is migratory and undergoes constant diffusion, while the other proliferates in an oxygen-dependent manner. The local oxygen concentration governs transitions between these phenotypes. We then ask whether the time scale of cyclic hypoxia, where oxygen is spatially homogeneous but fluctuates between normoxic and hypoxic levels over time, alters the dynamics of the reduced generalist model in the fast phenotype-switching regime compared with the full go-or-grow model. We observe that the reduced generalist phenotype model fails to capture the go-or-grow dynamics under rapid oxygen fluctuations. Our results predict that the tumor growth rate increases in the generalist-phenotype model under rapid oxygen fluctuations compared with tumors of the proposed go-or-grow model. We then explore whether oxygen consumption, which leads to spatial heterogeneity in oxygen concentration, plays a role in distinguishing the dynamics of the reduced model from the proposed go-or-grow model. We demonstrate that the tumor in the proposed go-or-grow model exhibits a higher growth rate than the tumor in the reduced generalist-phenotype model under a slow oxygen consumption rate.

## 1. Introduction

The go-or-grow dichotomy is a well-established mathematical modeling hypothesis whereby cell populations either migrate or proliferate, but not both simultaneously. This modeling framework has been used to describe many biological and medical conditions, mainly the tumor invasion process [1, 2, 3, 4, 5]. Under the go-or-grow hypothesis, cells can alternate between proliferative and migratory states via phenotypic switching, as observed in experiments [6]. This phenotypic switching may depend upon many factors, such as cell density and local oxygen levels [1, 7, 8]. Local oxygen levels in the tumor microenvironment may fluctuate between normoxic (this environment promotes migratory to proliferative phenotype switching) and hypoxic levels (this environment encourages proliferative to migratory phenotypic transition) [9], which is known as cyclic hypoxia [8, 10]. Moreover, oxygen in the tumor microenvironment is spatially heterogeneous as tumor cells consume oxygen, leading to the formation of an oxygen gradient from the tumor periphery to the core area [11, 12]. Therefore, oxygen concentration is spatio-temporally heterogeneous in the tumor microenvironment.

In many previous studies, phenotypic switching was associated with the local total tumor cell density and extracellular matrix (ECM) density [3, 7, 13]. That is, switching was governed by internal factors, allowing the go-or-grow model to be reduced to a generalist-phenotype cellular model, which effectively captured the go-or-grow dynamics in the fast phenotypic switching regime. However, phenotypic switching is also driven by the local oxygen level, an external factor. Alfonso et al. [14] derived a generalist-phenotype model from the go-or-grow model, assuming that phenotypic switching is governed by the local oxygen concentration in the fast phenotype-switching regime. However, our understanding of how the reduced generalist-phenotype model compares with the full go-or-grow model under fluctuating oxygen environments remains limited. In particular, oxygen fluctuations can occur over a wide range of time scales [10], and it remains unclear how the time scale of oxygen fluctuations affects the behavior of the generalist-phenotype and go-or-grow models. Furthermore, whether the timescale of oxygen consumption results in the generalist-phenotype model exhibiting faster speed of invasion as compared to the full go-or-grow model remains unexplored.

In this work, we propose a minimal go-or-grow model in spatio-temporal oxygen-heterogeneous environments using a system of reaction-diffusion equations. The proposed model consists of two equations for proliferative and migratory cells: migratory cells move via linear diffusion; proliferative cells divide based on local oxygen concentration; phenotypic switching is a function of local oxygen level; and an oxygen reaction-diffusion equation. The goal of this work is to capture go-or-grow dynamics through a single generalist phenotype equation model instead of the proposed two equation model for heterogeneous cell phenotypes. The model has three timescales: phenotypic switching, cell proliferation and migration, and oxygenation (such as diffusion, fluctuations, and consumption). In the fast phenotypic switching regime, we show that the model involving distinct phenotypic populations can be reduced to a model that consists of a single generalist-phenotype population, in which the diffusion coefficient and proliferation rate are linked to the phenotypic switching rates and proliferation rate of the two-population model. Our numerical results confirm that the single generalist-phenotype cell model behaviour is the same as that of the proposed go-or-grow model with heterogeneous populations for a specific choice of biologically meaningful phenotypic switching functions in both spatially and temporally heterogeneous oxygen environments. In the rapid oxygen fluctuation regime, where the timescale of oxygen fluctuations is very much shorter than those of phenotype switching and proliferation/movement, we observe, however, that the single-generalist-phenotype equation fails to reproduce the two-equation go-or-grow model behaviour. In addition, it is noted that the generalist phenotype cell tumor shows more aggressive invasion dynamics in the rapid oxygen fluctuation regime. Furthermore, we find that the reduced generalist-phenotype model fails to capture the dynamics of the go-or-grow model under a slow oxygen consumption regime. It is also noted that the generalist-phenotype model exhibits less aggressive invasion behavior than the full model under the slow oxygen consumption regime.

## 2. A two population go-or-grow model

Let *m*(**x**, *t*) and *p*(**x**, *t*) denote the densities of migratory and proliferative cells, respectively, at time *t* and position **x** = (*x, y, z*) in cartesian coordinate space and define the total cell density by

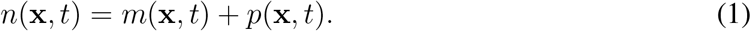

A minimal go-or-grow model, where phenotypic switching and proliferation depend on oxygen concentration (*c*(**x**, *t*)), is given by

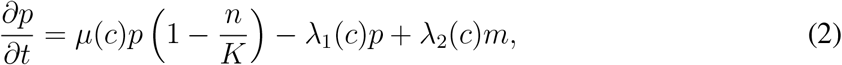

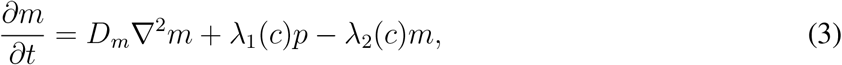

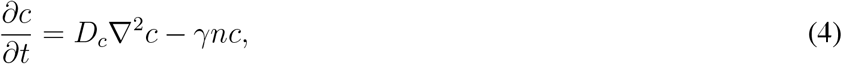

where *n* = *m* + *p* is the total cell density, 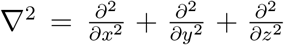. Here, the first term on the right-hand side of Eq. (2) describes the proliferative cells that proliferate logistically with linear growth rate *µ*(*c*), which is an increasing function of *c*, and have a constant carrying capacity, *K*. Proliferative cells adopt a migratory phenotype with a rate *λ*_1_(*c*), which is a decreasing of *c*. The term involving *λ*_2_(*c*) denotes migratory cells switching to proliferation and is an increasing function of *c*. Eq. (3) describes the diffusion (with constant diffusion coefficient *D*_*m*_, independent of *c*) of the migratory cells together with the corresponding switching terms. Here, and unless stated otherwise, we keep *µ*(*c*), *λ*_1_(*c*), and *λ*_2_(*c*) general, assuming them to be strictly non-negative functions of the oxygen concentration *c*. Eq. (4) describes the spatio-temporal dynamics of oxygen, where oxygen is assumed to diffuse, with constant coefficient of diffusion *D*_*c*_, and be consumed by both proliferative and migratory cells at the same rate *γ >* 0.

### 2.1. Reduction to single-equation for the coupled phenotypes model

Experimental studies reported that the time-scale of switching for both proliferative to migratory and migratory to proliferative phenotypes can be faster than that of cellular-level dynamics, such as cell proliferation and cell migration [15]. We define the parameter *ε* to be the ratio between the phenotypic switching time scale and the population dynamics time scale, namely

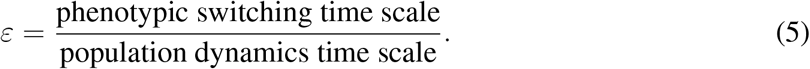

The condition 0 *< ε ≪* 1 implies that phenotypic adaptation occurs much faster than migration or proliferation. Fast phenotypic switching is modeled for Eqs. (2)-(3) by assuming [7]

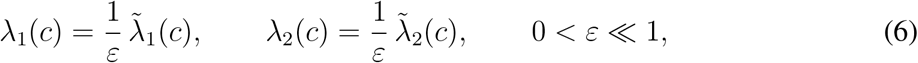

where 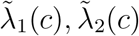 are *O*(1).

Hence, the typical fast-switching system takes the form

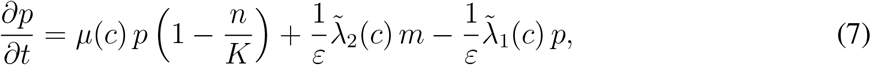

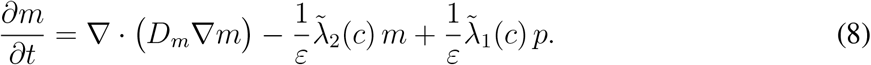

In the limit *ε →* 0, the leading-order balance yields the relation

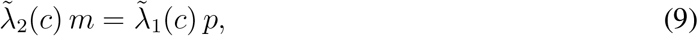

and the two cell densities *p* and *m* can be related to the total cell density (*n*) via

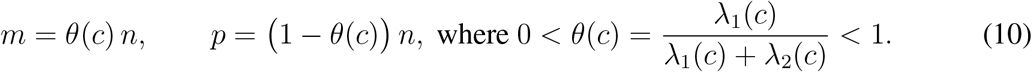

Substituting these expressions into the total density equation (sum of Eqs. (7) and (8)) leads to the reduced two-equation model

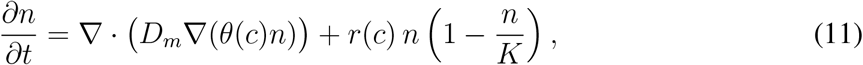

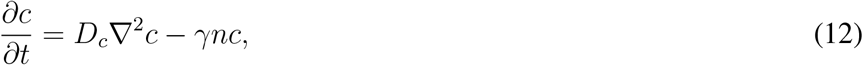

with effective oxygen-dependent proliferation rate

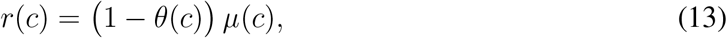

and the generalist phenotype cell’s diffusivity depends on oxygen concentration.

## 3. Numerical results

In this section, we explore numerical results for the choice of a specific phenotype switching function. We restrict our study to the spatially one-dimensional case, and take the functional forms of the proliferation rate and phenotype switching rates to be, respectively,

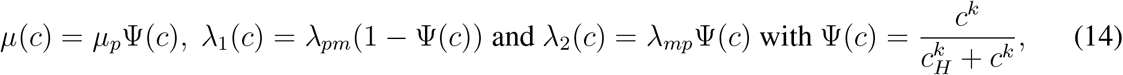

where *µ*_*p*_, *λ*_*pm*_, and *λ*_*mp*_ represent the rates of cell proliferation, the transition from a proliferative to a migratory phenotype, and the transition from a migratory to a proliferative phenotype, respectively. Here Ψ(*c*) is a Hill function. This choice satisfies the conditions that both *µ*(*c*) and *λ*_2_(*c*) are increasing functions of *c*, and that *λ*_1_(*c*) is a decreasing function of *c*, so, 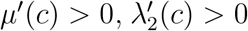 and 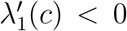, where ^*′*^ denotes derivative with respect to *c*. Additionally, *c*_*H*_ *>* 0 denotes the hypoxic threshold value of oxygen concentration, and *k >* 0 refers to the Hill coefficient. Hence,

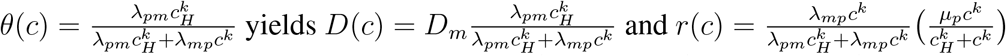

To obtain numerical results, we employed the forward time central difference in space (FTCS) scheme within a finite difference framework in a one-dimensional Cartesian domain [0, *L*] with tumor center at *x* = 0. We assume that migratory cells do not leave the domain. Hence, for the heterogeneous model (Eqs. (2) and (3)) we impose no-flux boundary conditions for Eq. (3) at *x* = 0 (as a symmetry condition) and *L*. For the oxygen equation (Eqs. (4) and (12)), we assume that the oxygen level is a fixed positive constant at the outward boundary, i.e., *c*(*L, t*) = *c*_*∞*_ *>* 0. Due to symmetry, we impose the boundary condition, 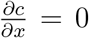 at *x* = 0. Also, we assumed that the tumor initially grows from proliferative cells only, and the initial conditions are considered as *p*(*x*, 0) = 0.2*ℋ* (*x −* 2) and *m*(*x*, 0) = 0 for *x ∈* [0, *L*] with *L* = 10. For the generalist phenotype model (Eq. 11), we use the no-flux condition at the boundary of the domain [0, *L*] and initial condition as *n*(*x*, 0) = 0.2*ℋ* (*x −* 2) for *x ∈* [0, *L*]. The details of the numerical method can be found in the Numerical method section of Supplementary file.

### 3.1. During cyclic hypoxia

First, we begin with the case where we assume that oxygen is spatially homogeneous but temporally varying. In most in vitro experiments done so far to recapitulate cyclic hypoxia, spatial heterogeneity has not been the focus, instead temporal fluctuations of oxygen levels have been imposed on cells [8, 16]. Hence, oxygen is spatially homogeneous (i.e., *c*(**x**, *t*) = *c*(*t*)), and *c*(*t*) fluctuates over time between oxygen-rich and oxygen-poor levels. Therefore, cyclic hypoxia can be modeled as

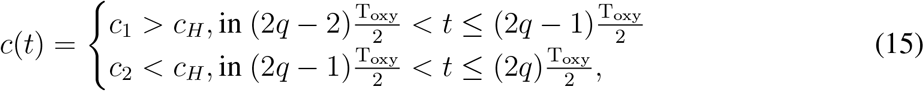

where *T*_oxy_ is the oxygen fluctuation period and *q* = 1, …, *s* with *sT*_oxy_ being the total time of interest. The corresponding reduced equation Eq. (11) becomes

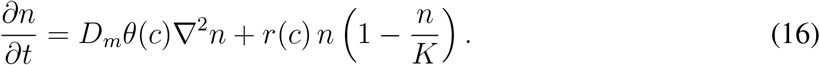

In this problem, we have three timescales, namely 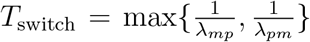 denotes the phenotypic adaptation time scale, 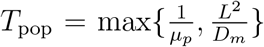 the population dynamics time scale associated with diffusion and proliferation, and oxygen fluctuation timescale (*T*_oxy_). In the fast phenotype switching regime i.e., *T*_switch_ *≪ T*_oxy_, we observed that the total density of the distinct heterogeneous population model follows the same time course evolution as for the generalist phenotype heterogeneous model (Figure 1A). Under this condition, phenotypic switching equilibrates before significant migration or growth occurs. However, in the rapid oxygen oscillation regime i.e., *T*_oxy_ *≪ T*_switch_ *< T*_pop_, cells are unable to adapt to instantaneous oxygen variations. As a result, rapid periodic oxygen fluctuations do not induce go-or-grow alternation in this regime (Figure 1B). Figure 1B(ii) shows that the spatial position of the tumor with generalist-phenotype cells is ahead of that of specialist-phenotype tumor cells for the rapid oxygen fluctuation regime. This suggests that tumors with generalist-phenotype cells are more aggressive than tumors composed of specialist tumor cells under rapid oxygen fluctuations.

**Figure 1:**
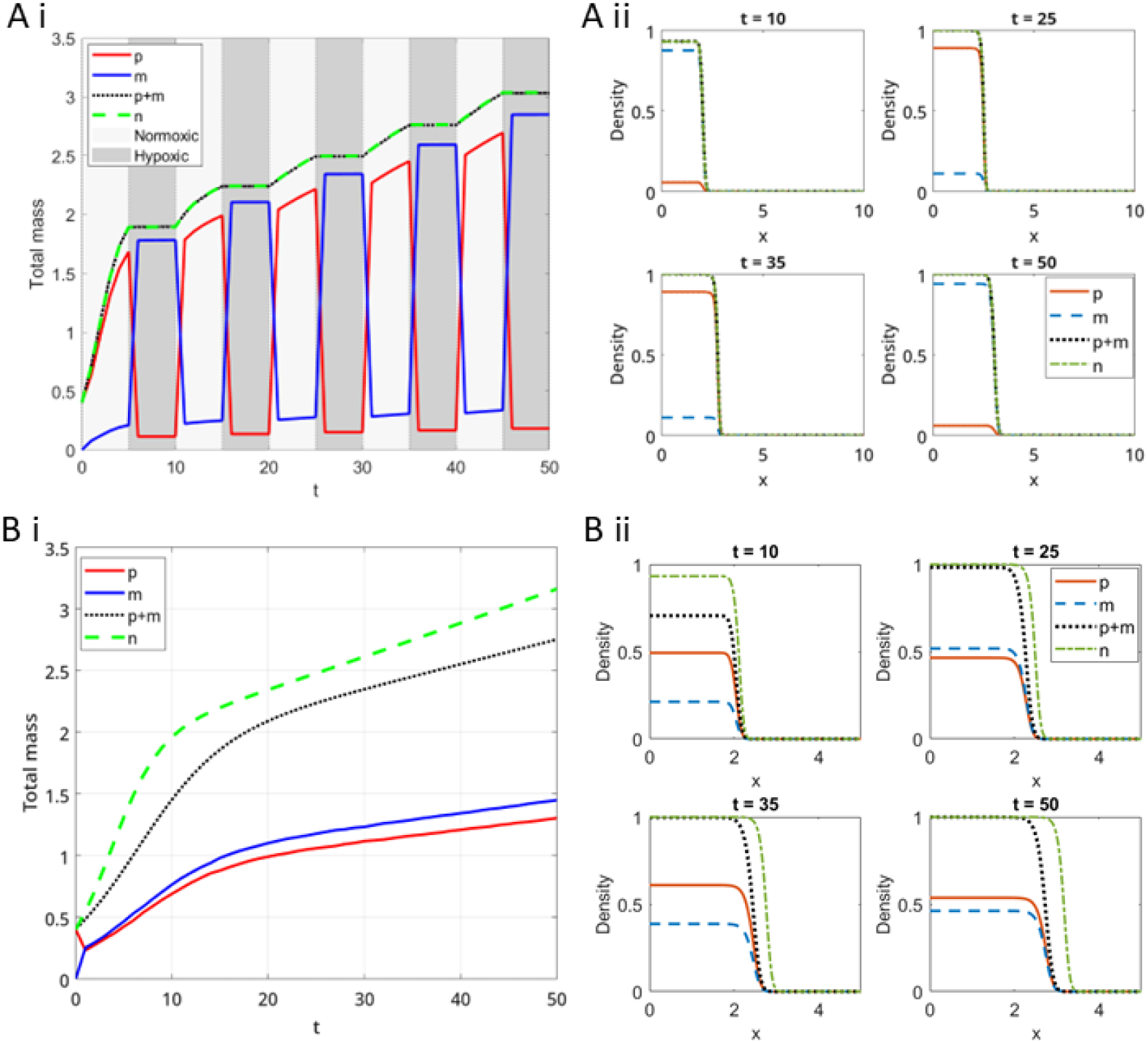
Time evolution of the proliferative cell density (*p*), migratory cell density (*m*), total (sum of proliferative and migratory cell density) and generalist phenotype (*n*) cell density in (A) a fast phenotype regime, (B) a rapid oxygen oscillation regime for the case where oxygen is spatially homogeneous but cycles in time between normoxic and hypoxic levels. Here, *D*_*m*_ = 0.01, *λ*_*pm*_ = 10, *λ*_*mp*_ = 20, *µ*_*p*_ = 1, *L* = 10, *K* = 1, *c*_*H*_ = 0.5 and the Hill coefficient *k* = 3. The period of cyclic hypoxia (*T*_oxy_) is 10 in (A) and 0.001 in (B).

### 3.2 Spatially heterogeneous oxygen environments

Spatial oxygen heterogeneity arises from oxygen consumption by tumor cells and the diffusion distance from the oxygen source [12, 17]. Experimental studies have reported that the variability in oxygen diffusivity across different cancers is insignificant, whereas the oxygen consumption rate can vary substantially and depends on the cell type and tumor location [18]. In this section, we investigate whether the reduced generalist phenotype model captures the go-or-grow model under fast phenotypic-switching regimes across different oxygen consumption rates.

In this case, the oxygen diffusion equation (4) is solved with a symmetric boundary condition at *x* = 0 (i.e. zero flux, under the assumption that our domain is capturing the dynamics in half of a tumor centered at *x* = 0) and a Dirichlet boundary condition at *x* = *L*, such that *c*(*L, t*) = *c*_*∞*_ *>* 0 coupled with the specialist cellular model (Eqs. (2) and (3)). Simulation results show that the total cell density of the proposed go-or-grow model has the same time behavior as the reduced generalist phenotype model (Eqs. (11) and (12)) in the fast phenotypic switching regime when the oxygen consumption rate is high (Figure 2A i). It is noted that the total density initially increases and then reaches a steady state. This occurs because, during the initial phase, the oxygen level in the tumor region is sufficiently high, allowing proliferative cells to proliferate rapidly and thereby increase the total tumor density. However, as time progresses, an oxygen gradient develops within the tumor region, and the oxygen level falls below the threshold value *c*_*H*_ (Figure 2A ii). Consequently, proliferative cells switch to a migratory phenotype, and their proliferation rate is reduced. In addition, we found that the reduced generalist-phenotype model is unable to capture the go-or-grow dynamics when oxygen consumption by tumor cells is sufficiently low. In this regime, the spatial position of the generalist-phenotype cells lags behind that of the total cell density of the go and grow phenotypes (Figure S1). From the reduced model, we observed that the flux of generalist-phenotype cells depends on the oxygen level at their spatial location, which has a local effect that decreases cell motility. In contrast, the cells in the go-or-grow model can move freely through space.

**Figure 2:**
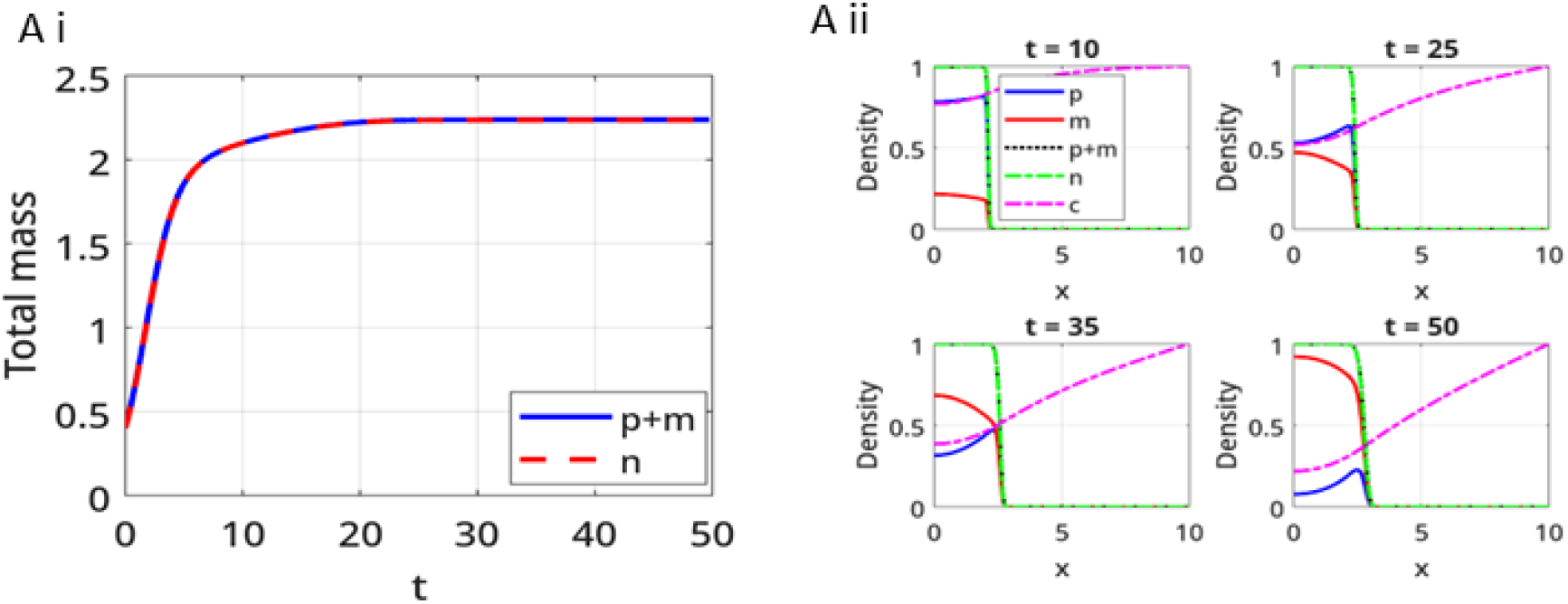
Time evolution of the proliferative (*p*), migratory (*m*) and total (sum of proliferative and migratory cells) and generalist phenotype (*n*) cell density in a spatially heterogeneous oxygen environments, where there is a source of oxygen at the right-hand boundary. Here, *D*_*m*_ = 0.01, *λ*_*pm*_ = 10,*λ*_*mp*_ = 20, *µ*_*p*_ = 1, *L* = 10, *K* = 1, *c*_*H*_ = 0.5 and the Hill coefficient *k* = 3. The oxygen diffusion coefficient *D*_*c*_ = 0.1 and *γ* = 0.1, *c*_*∞*_ = 1.

## 4. Model analysis for oxygen-rich and -deprived conditions

We simplify the switching function Ψ(*c*) to be (in the limit *k → ∞*) Ψ(*c*) = *ℋ* (*c − c*_*H*_), where *H*(.) is the Heaviside function:

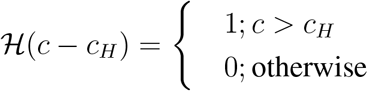

Hence, the rates for cell proliferation, the transition from proliferative to migratory, and the transition from migratory to proliferative for Eqs. (2)-(3) given in Eqs. (14), reduce to

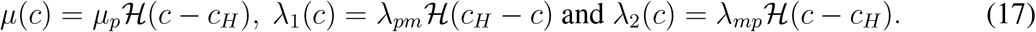

We now consider the case where oxygen levels are constant in time and spatially homogeneous and, as before, we assume that the tumor grows from proliferative cells only, so, initially, *p* = *p*_0_ and *m* = 0, hence *n* = *p*_0_.

**For oxygen-rich conditions, i.e**., *c > c*_*H*_ : In this case, proliferative cells cannot transition into migratory cells so, from the initial conditions, the migratory cell density remains at zero. As a result, the go-or-grow model reduces to:

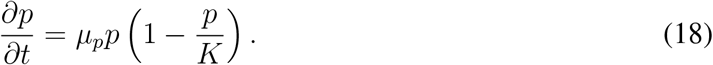

Here *θ*(*c*) = 0, so, *D*(*c*) = 0 and *r*(*c*) = *µ*_*p*_. Hence, the single generalist equation (11) becomes

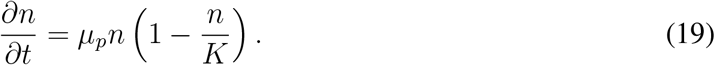

Clearly, Eq. (18) and Eq. (19) are identical.

**For oxygen-deprived conditions, i.e**., *c < c*_*H*_ : The go-or-grow model becomes

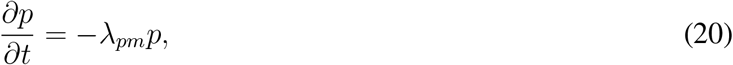

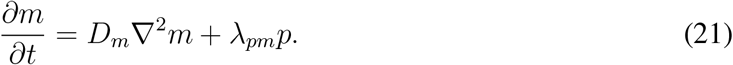

By adding Eqs. (20) and (21), we have

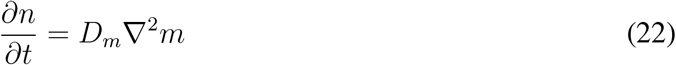

However, under this oxygen-deprived condition, all proliferating cells (p) switch to migratory cells (m) rapidly, as we are in the fast phenotypic switching regime (Figure S2). Hence, *m* is equivalent to *p* + *m* i.e., *n* after initial transient dynamics.

In this case, *θ*(*c*) = 1, so, *D*(*c*) = *D*_*m*_ and *r*(*c*) = 0. Therefore, the single generalist equation (11) becomes

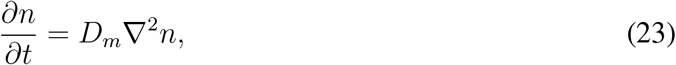

which is exactly same as the sum of Eqs. (20) and (21) in fast phenotype switching regime, which has observed in Figure S2 v-vi. However, we found that the reduced generalist-phenotype model fails to capture the dynamics of the specialist-phenotype go-or-grow model at timescales shorter than the phenotypic switching timescale (zoomed sections of Figure S2 i-iv).

## 5. Discussion and future direction

In this work, we analyzed a minimal go-or-grow model, in which cells can either proliferate via logistic growth or move via constant diffusion, with oxygen-dependent proliferation rate and phenotypic switching rates. We derived a similar relationship between the model of two phenotypic cell types (“specialists”) and a model for a single generalist in the fast-phenotype-switching regime with oxygen concentration-dependent phenotypic switching, which is shown in [14, 3]. This relation suggests that the diffusion coefficient and proliferation rate in the generalist model are determined by oxygen-dependent switching rates between specialist cells that are phenotypically distinct in the go-or-grow model. Tumors experience oxygen fluctuations in the vicinity of blood vessels and during ongoing angiogenesis. However, our understanding is limited to whether this reduced generalist phenotype model captures the go-or-grow dynamics in oxygen fluctuation regimes. Moreover, the oxygen consumption rate varies across tumors, depending on tumor location and cell type [19, 20]. This work also investigated whether the oxygen consumption rate confers any benefit to generalist cells during invasion, compared with tumors of specialist cells.

Our simulation results show that in the fast phenotype-switching regime, the generalist-phenotype model closely resembles a two-phenotype population model in spatio-temporal heterogeneous oxygen settings. We have also shown that if the oxygen oscillation timescale is very fast (for example, near blood vessels), then the generalist-phenotype (“generalist”) model fails to accurately capture the go-or-grow hypothesis. It showed that the generalist phenotype tumor demonstrates higher invasion than the tumors of the full go-or-grow model in a rapid oxygen fluctuation environment. In addition, we observed that the reduced generalist-phenotype tumor model exhibits the same dynamics as the go-or-grow model when the oxygen consumption rate is high. We found that generalist-phenotype tumors show higher invasiveness when the oxygen consumption rate is low, compared to the specialist go-or-grow model. A similar comparison was carried out in a previous study [1], which showed that the invasion speeds could swap depending on parameter values. However, they observed this behavior using two distinct models for the generalist and specialist phenotypes of cells. In contrast, in our work, the model for the generalist cell type has been obtained by reducing the proposed model for the specialist cell type. Furthermore, in this work, we have only considered linear diffusion of migratory cells, however, more complex models have been proposed in which migration is affected by total cell density and the extracellular matrix [1], giving, potentially, a more realistic picture of tumor invasion.

## Data availability

The code for numerical simulations are available at GitHub in the following link: https://github.com/gsadhuIITG/Single_Eqns.

## Acknowledgment

GS and MKJ are supported by Param Hansa Philanthropies. PKM would like to thank the Royal Society Yusuf Hamied Visiting Fellowship scheme which funded his visit to IISc Bangalore. For the purpose of open access, the author has applied a CC BY public copyright license to any author accepted manuscript arising from this submission.

## Conflict of interest

The authors declare no conflict of interest.

## Supplementary Information

### Numerical method

The domain [0, *L*] is divided into *N* equal grid spaces Δ*x*, with 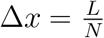, and Δ*t* is the time-step. We define 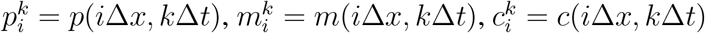 and 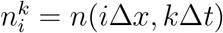. We employ forward difference scheme for the temporal derivative and central difference for the spatial derivative to discretize the governing equations.

The discretized form of Eq. (2) becomes

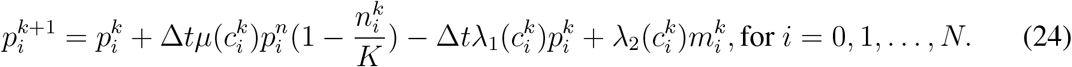

The discretized form of Eq. (3) becomes

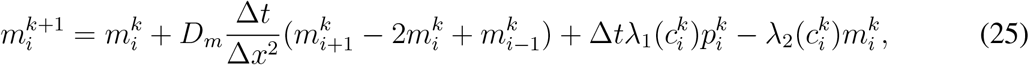

for *i* = 1, 2, …, *N* 1.

At boundary points i.e., *i* = 0, *N*, we use ghost point criteria and the corresponding discretized form is prescribed as

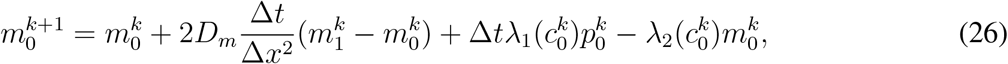

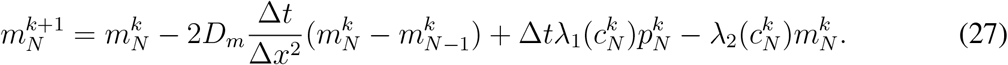

The oxygen diffusion equation (4) discretize as

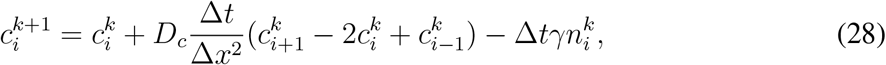

for *i* = 1, 2, …, *N* 1. At *i* = 0, we use ghost point criteria and the corresponding discretized form is written as

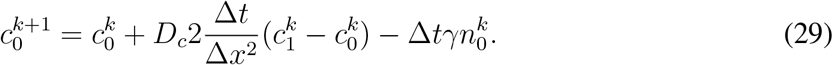

The discretized form of Eq. (11) becomes

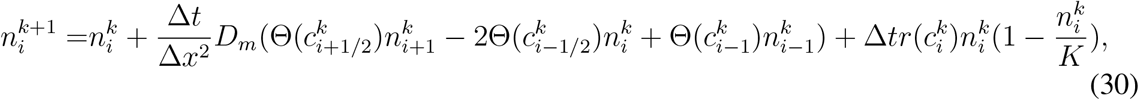

for *i* = 1, 2, …, *N −* 1. At the boundary points *i* = 0 and *N*, it takes the form

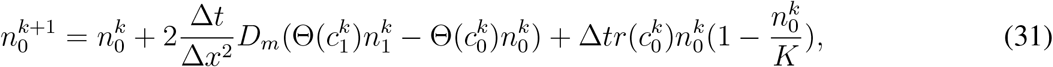

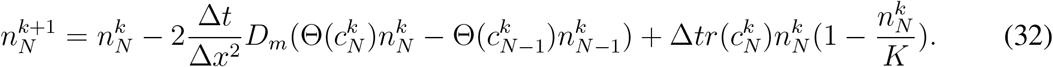

Numerical stability is ensured through the choice of time step length 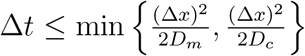 in each time step.

**Figure S1:**
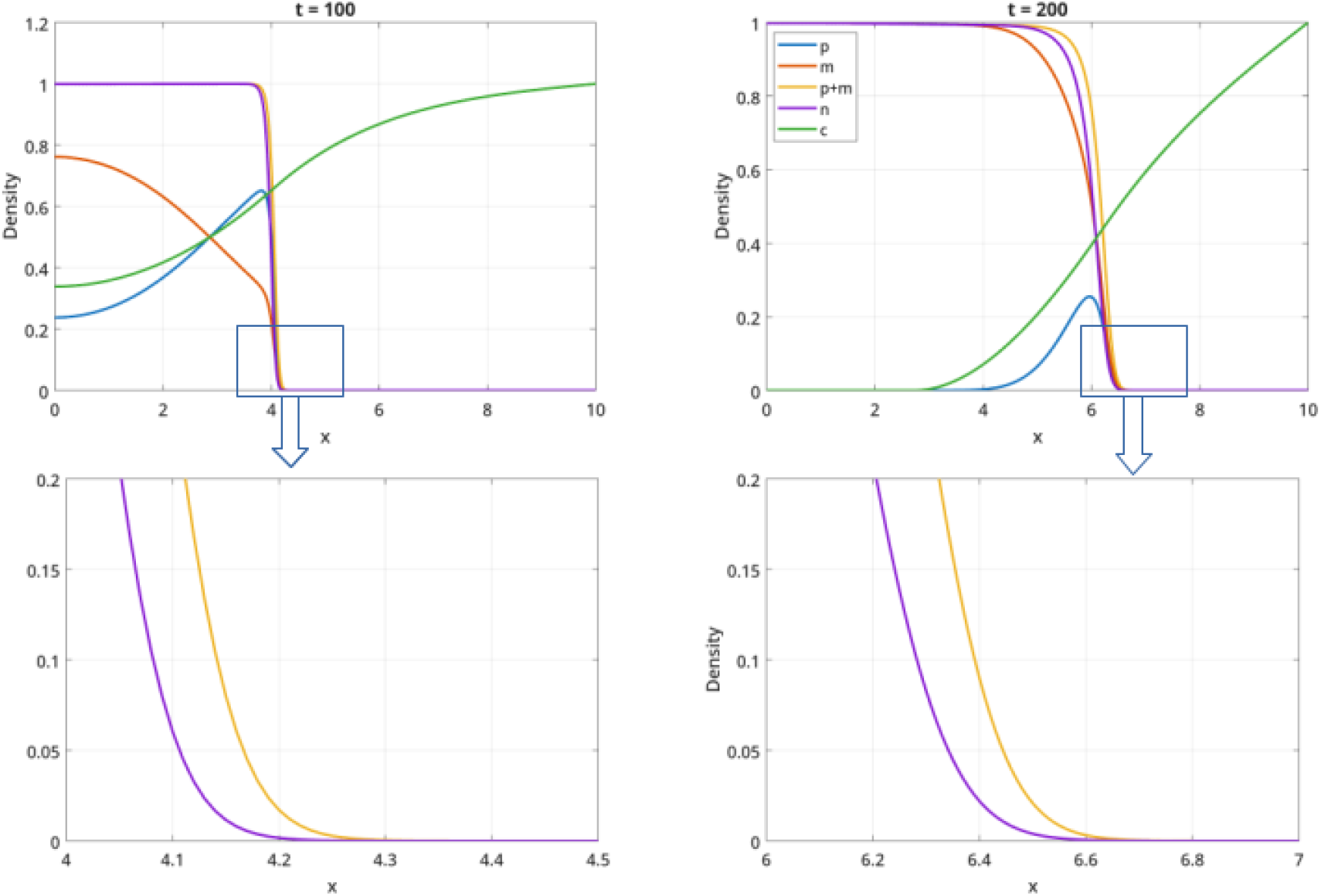
The spatial distribution of *p, m, p* + *m, n* and *c* at *t* = 100 and 200. In the zoomed-in portion, *p* + *m* and *n* density distribution over the spatial domain at *t* = 100 and 200.

**Figure S2:**
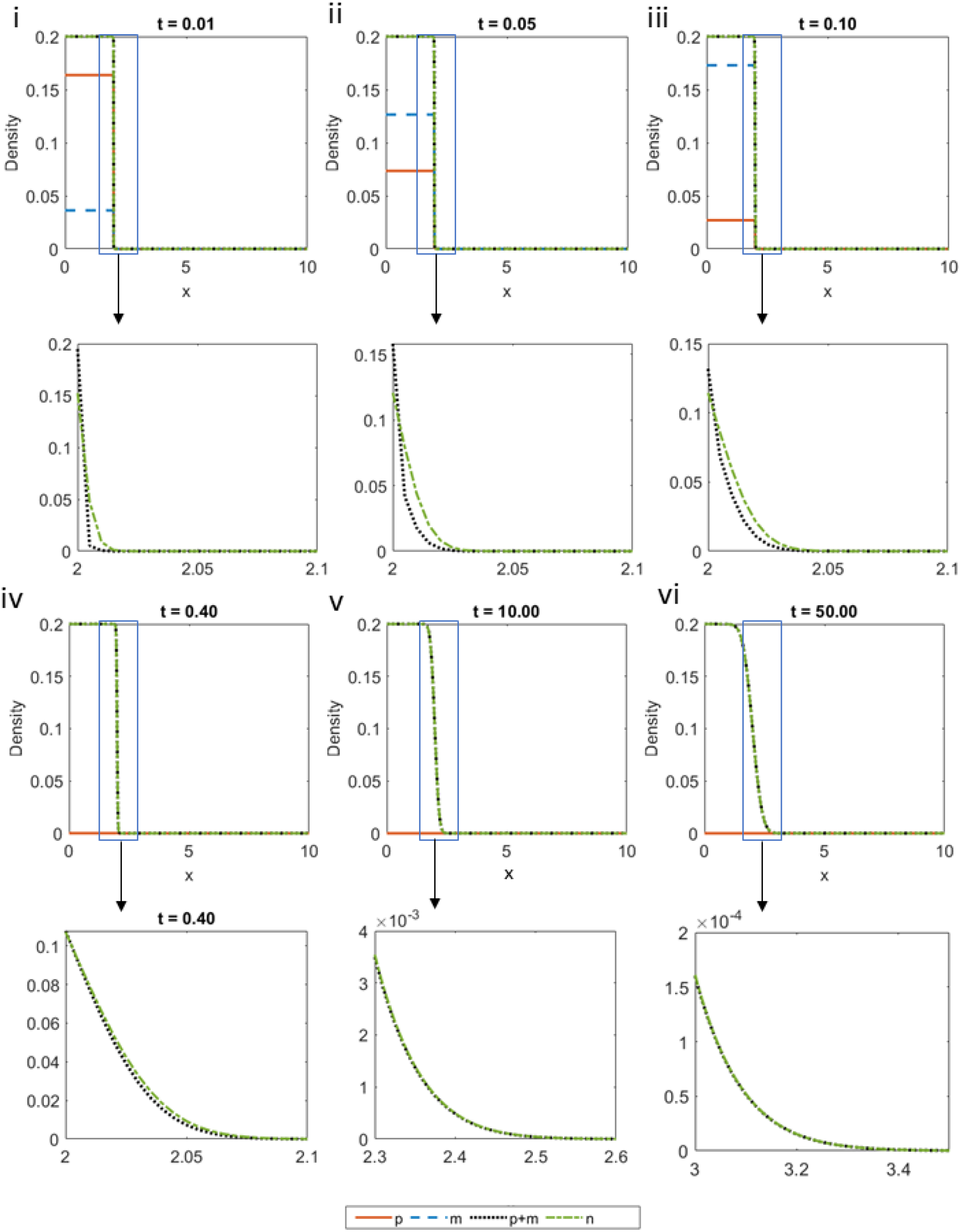
The spatial distribution of *p, m, p* + *m* and *n* at *t* = 0.01, 0.05, 0.1, 0.4, 10 and 50. In the zoomed-in portion, *p* + *m* and *n* density distribution over the spatial domain.

## Notes

### Competing Interest Statement

The authors have declared no competing interest.

